# Dynamic aqueous multiphase reaction system for simple, sensitive and quantitative one-pot CRISPR-Cas12a based molecular diagnosis

**DOI:** 10.1101/2020.03.21.001644

**Authors:** Kun Yin, Xiong Ding, Ziyue Li, Hui Zhao, Kumarasen Cooper, Changchun Liu

## Abstract

Recently, CRISPR-Cas technology has opened a new era of nucleic acid-based molecular diagnostics. However, current CRISPR-Cas-based nucleic acid biosensing has largely a lack of the quantitative detection ability and typically requires separate manual operations. Herein, we reported a dynamic aqueous multiphase reaction (DAMR) system for simple, sensitive and quantitative one-pot CRISPR-Cas12a based molecular diagnosis by taking advantage of density difference of sucrose concentration. In the DAMR system, recombinase polymerase amplification (RPA) and CRISPR-Cas12a derived fluorescent detection occurred in spatially separated but connected aqueous phases. Our DAMR system was utilized to quantitatively detect human papillomavirus (HPV) 16 and 18 DNAs with sensitivities of 10 and 100 copies within less than one hour. Multiplex detection of HPV16/18 in clinical human swab samples were successfully achieved in the DAMR system using 3D-printed microfluidic device. Furthermore, we demonstrated that target DNA in real human plasma samples can be directly amplified and detected in the DAMR system without complicated sample pre-treatment. As demonstrated, the DAMR system has shown great potential for development of next-generation point-of-care molecular diagnostics.

## Introduction

Molecular diagnostics is critical for the identification of pathogens and genotyping, which makes an outstanding contribution to clinical diagnostics, biosecurity and environmental monitoring.^1^, Nucleic acid amplification testing, such as polymerase chain reaction (PCR), is the most commonly used technique in molecular diagnostics.^1^ However, PCR technique typically requires sophisticated system and well-trained operator.^2^ Therefore, there is an unmet need to develop a novel nucleic acid-based molecuar testing method for simple, rapid, sensitive, reliable and cost-effective detection at the point of care.

Nucleic acid isothermal amplification detection,^3, 4^ such as recombinase polymerase amplification (RPA), loop mediated isothermal amplification (LAMP), is an attractive alternative to conventional PCR method because of its outstanding advantages including simple, rapid and low cost. However, there remains a challenge to adapt it to develop an accurate and reliable point of care (POC) diagnostics for clinical applications due to undesired non-specific signals (e.g., false-positive).^5^ Recently, CRISPR-Cas system, has been widely applied as a revolutionary gene-editing technique in epigenetic engineering, gene regulation and genetic screening.^6^ Besides its extraordinary gene editing ability, it shows great promise for the next-generation of rapid and highly sensitive nucleic acid detection. Recently, a series of Cas effectors including Cas9, Cas12a and Cas13 have been developed to establish CRISPR-Cas-based nucleic acid biosensing detection.^7–10^ For example, the Cas12a owns collateral cleavage activities on single stranded DNA (ssDNA).^8^ The cleavage activity of Cas12a can be activated, and indiscriminately cleave the collateral ssDNA reporter once recognizing their specific DNA targets.^9^ To achieve a high sensitive molecular detection, it is necessary to combine target nucleic acid amplification (e.g., RPA) with CRISPR-based detection. For instance, DETECTR method has recently been developed for highly sensitive and specific nucleic acid detection by combining the RPA amplification with Cas12a detection.^8^ However, due to poor biocompatibility of two different reaction systems, the DETECTR assay typically requires separate target amplification and detection steps. Such “two-step” assay has some drawbacks for POC diagnostics because: i) transfering of RPA amplicons exposes the nucleic acid-rich sample (e.g., RPA amplicons) to the environment, potentially increasing the risk of carry-over contamination, and ii) it cannot accurately quantify the target nucleic acids due to the separate target amplification.^8, 11–13^

## Results

### Design of dynamic aqueous multiphase reaction system

Herein, we have developed a dynamic aqueous multiphase reaction (DAMR) system for simple, rapid, sensitive and quantitative detection of nucleic acids, where the RPA reaction and CRISPR-Cas12a detection were carried out in spatially separated but connected phases in one-pot. The DAMR system was established by taking advantage of density difference of sucrose concentration (inset of Fig. 1 and Fig. S1). This miscible multiphase system provides a unique dynamic diffusion interface, which can combine incompatible but related reactions together and enable one-pot RPA-CRISPR/Cas12a molecular detection. As shown in Fig. 1, target nucleic acids are firstly amplified by the RPA reaction in high-density bottom phase. Next, the amplified nucleic acids dynamically diffuse to the low-density top-phase and sequence-specifically trigger the nonspecific cleavage activity of Cas12a endonuclease with crRNA, which further cleaves fluorophore quencher (FQ)-labeled ssDNA probe to generate fluorescent signal for nucleic acid detection. To our best knowledge, it is the first time to adapt a dynamic aqueous multiphase system to achieve incompatible biochemical reactions in one-pot.

**Figure 1.**
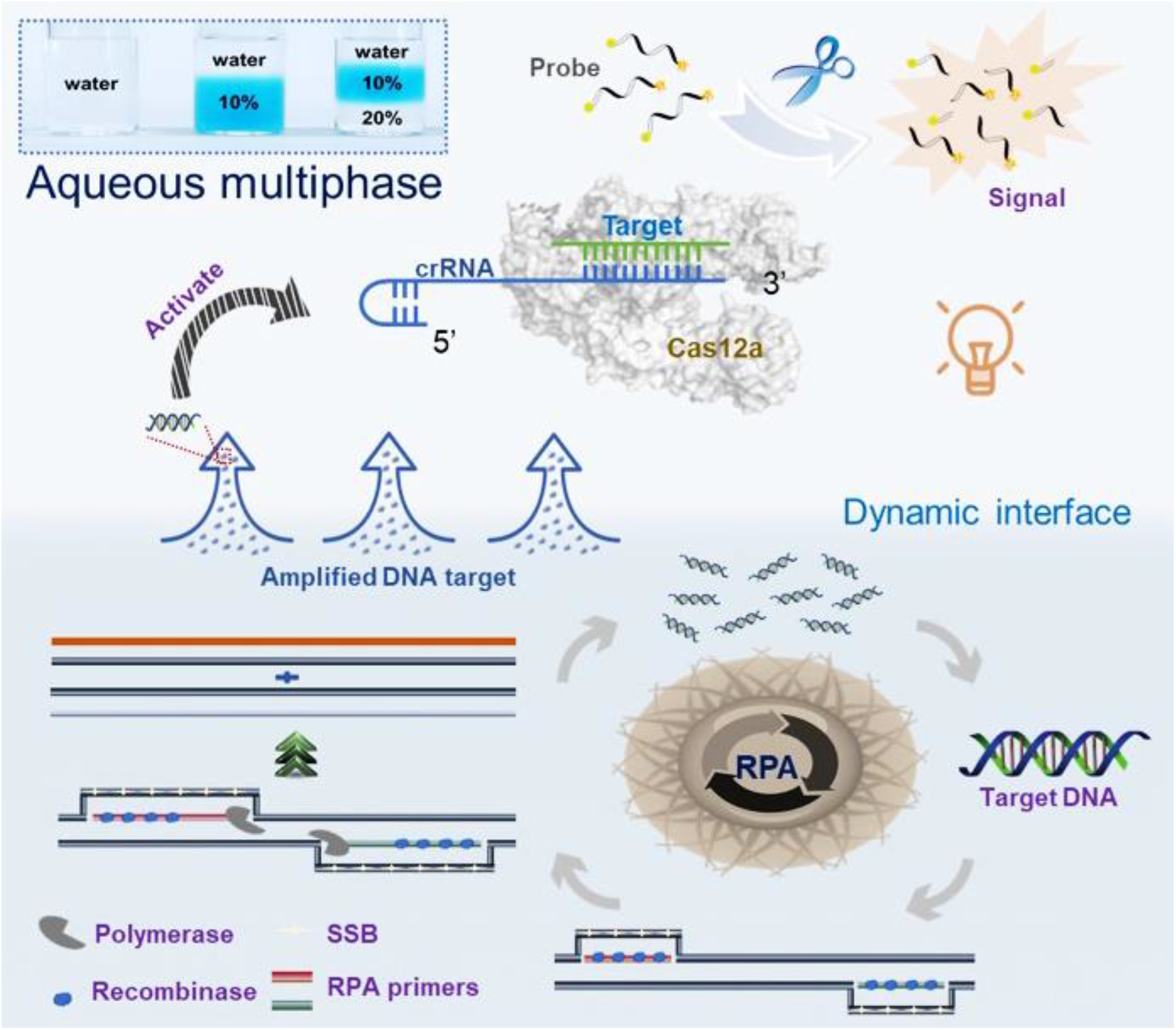
Schematic illustration of dynamic aqueous multiphase reaction (DAMR) system for simple, sensitive and quantitative DNA detection by one-pot RPA/CRISPR-Cas12a assay. Inset is an image of aqueous multiphase system with different sucrose concentrations.

To evaluate the performance of the DAMR system, 10^4^ copies of human papillomavirus (HPV) 16 DNA was detected by using 20 μL RPA reaction solution with 10% sucrose as the bottom phase and 10 μL CRISPR-Cas12a fluorescence detection solution as the top phase. For comparison, we thoroughly mixed all-above mentioned reaction reagents in a one-phase system for HPV16 testing. As shown in Fig. 2A, obvious fluorescent signals can be observed after 30-min incubation in the DAMR system, whereas not in the one-phase system. Interestingly, the fluorescent signal firstly generated at the top phase of CRISPR-Cas12a detection and then diffused into the bottom phase of the RPA reaction, which indicated that the DAMR system provided relatively independent, but dynamical diffusion environment during the incubation process. As shown in Fig. 2A and S2B, no fluorescent signal can be observed in the one-phase system until incubated at 37 °C for 48 hours, which is 100 times slower than that of the DAMR system. Additionally, the fluorescence signal in the DAMR system is more than 100 times higher than that of the one-phase reaction in the same incubation time (Fig. 2A-ii and S2), which may be attributed to the unique dynamic multiphase reactions in our DAMR system. On one hand, the bottom phase of the DAMR system provided an independent reaction environment for the RPA amplification due to density-driven phase separation and sucrose hydrogen bond network.^14, 15^ On the other hand, the small nucleic acid amplicons generated in the bottom phase dynamically diffused into the top phase and triggered the cleavage activity of CRISPR-Cas12a for fluorescence detection.

**Figure 2.**
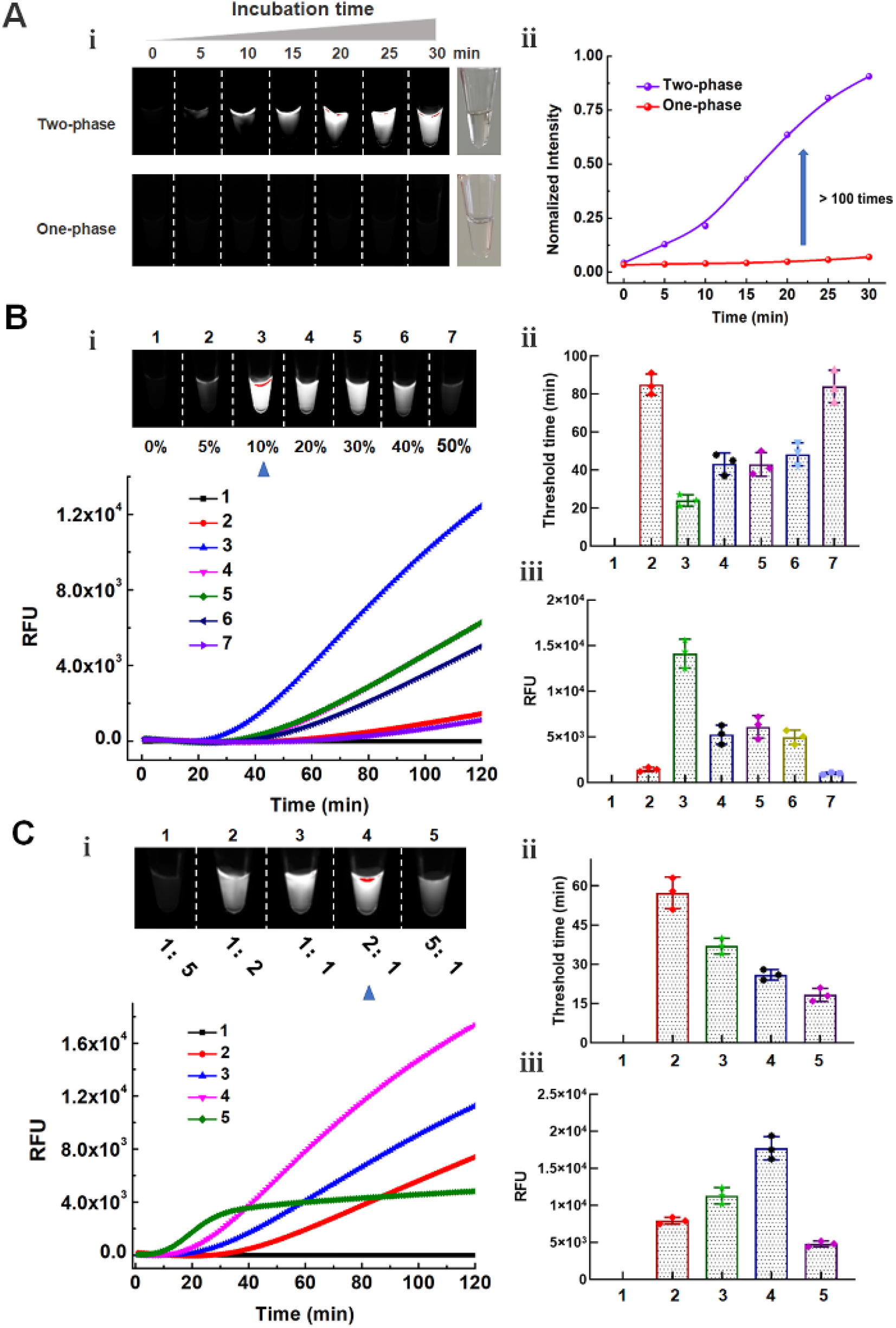
Optimization of one-pot RPA/CRISPR-Cas12a detection in the DAMR system. A: Fluorescent monitoring of the one-pot RPA/CRISPR-Cas12a reactions at different incubation times in the DAMR and one-phase system, respectively. B: Series sucrose concentration (0%, 5%, 10%, 20%, 30%, 40%, and 50%) were added to 20 μL RPA bottom phase and tested in the DAMR system. C: Effect of the volume ratios (1: 1:5, 1:2, 1:1, 2:1, 5:1) of the RPA bottom phase and CRISPR-Cas12a top phase on DNA detection in the DAMR system. Error bars denote s.d. (n=3).

### Optimization of dynamic aqueous multiphase reaction system

In our DAMR system, the dynamic diffusion between different phases plays a crucial role in connecting incompatible but related reaction systems, which is dependent on sucrose concentration and the volume ratio of two phases.^15, 16^ To optimize our DAMR system, we added different concentration sucrose (from 5% to 50%) into 20 μL RPA reaction solution with 10^4^ copies of HPV16 DNA target as the bottom phase and 10 μL CRISPR-Cas12a detection buffer as the top phase. The fluorescence signal during incubation was monitored in real-time by PCR machine and the endpoint fluorescent images were taken by ChemiDoc™ MP Imaging System. As shown in Fig. 2B, the RPA reaction solution with 10% sucrose exhibited the strongest fluorescent signal (Fig. 2B-i and 2B-iii) with the shortest threshold time (Fig. 2B-ii). Next, we optimized the volume ratio of the RPA bottom phase (10% sucrose) and CRISPR-Cas12a top phase. As shown in Fig. 2C, the fastest reaction could be achieved in the volume ratio of 5:1 but its fluorescent intensity was much lower than that of the volume ratio of 2:1. Considering the strong fluorescence signal plays a crucial role in visual detection of nucleic acid, 20 μL RPA reaction solution with 10% sucrose as the bottom phase and 10 μL CRISPR-Cas12a as the top phase were used in our optimized DAMR system for the one-pot RPA/CRISPR-Cas12a detection.

### Quantitative detection and sensitivity

Pathogens quantitative detection plays a crucial role in estimation of health risks and classification of disease severity.^17^ In the previous study, RPA amplification was separated from CRISPR-Cas12a detection due to poor compatibility of these two reaction systems.^8^

Therefore, the high concentration of DNA amplicons produced from the “first-step” RPA amplification leads to saturated fluorescence signals at the “second-step” regardless of initial nucleic acid target concentration, which could not realize accurate quantitative detection (Fig. S3 and S4). To investigate the quantitative detection performance of our DAMR system, tested ten-fold serial dilution of HPV 16 and 18 DNA by one-pot RPA/CRISPR-Cas12a method in the DAMR system, respectively. As shown in Fig. 3A-C and S5A-C, we can consistently detect 10 copies of HPV16 DNA and 100 copies of HPV18 within one hour, respectively, which was comparable with that of “two-step” method. In addition, a good linear relationship between the threshold time and DNA target concentration was achieved (Fig. 3D and Fig. S5D), which confirmed the excellent quantitative detection capability of our DAMR system. Furthermore, the endpoint fluorescence signal can be read directly by naked eyes under blue light (Fig. 3B), enabling simple, rapid and affordable point-of-care diagnosis without need for complicated equipment. Also, with specific primers during RPA reaction and specific crRNA during CRISPR-Cas12a based detection, satisfied selectivity was achieved using one-pot RPA/CRISPR-Cas12a in our DAMR system (Fig. 3E and Fig. S6). Therefore, our one-pot RPA/CRISPR-Cas12a assay in the DAMR system provides a simple, rapid, sensitive and reliable method for the quantitative detection of nucleic acids.

**Figure 3.**
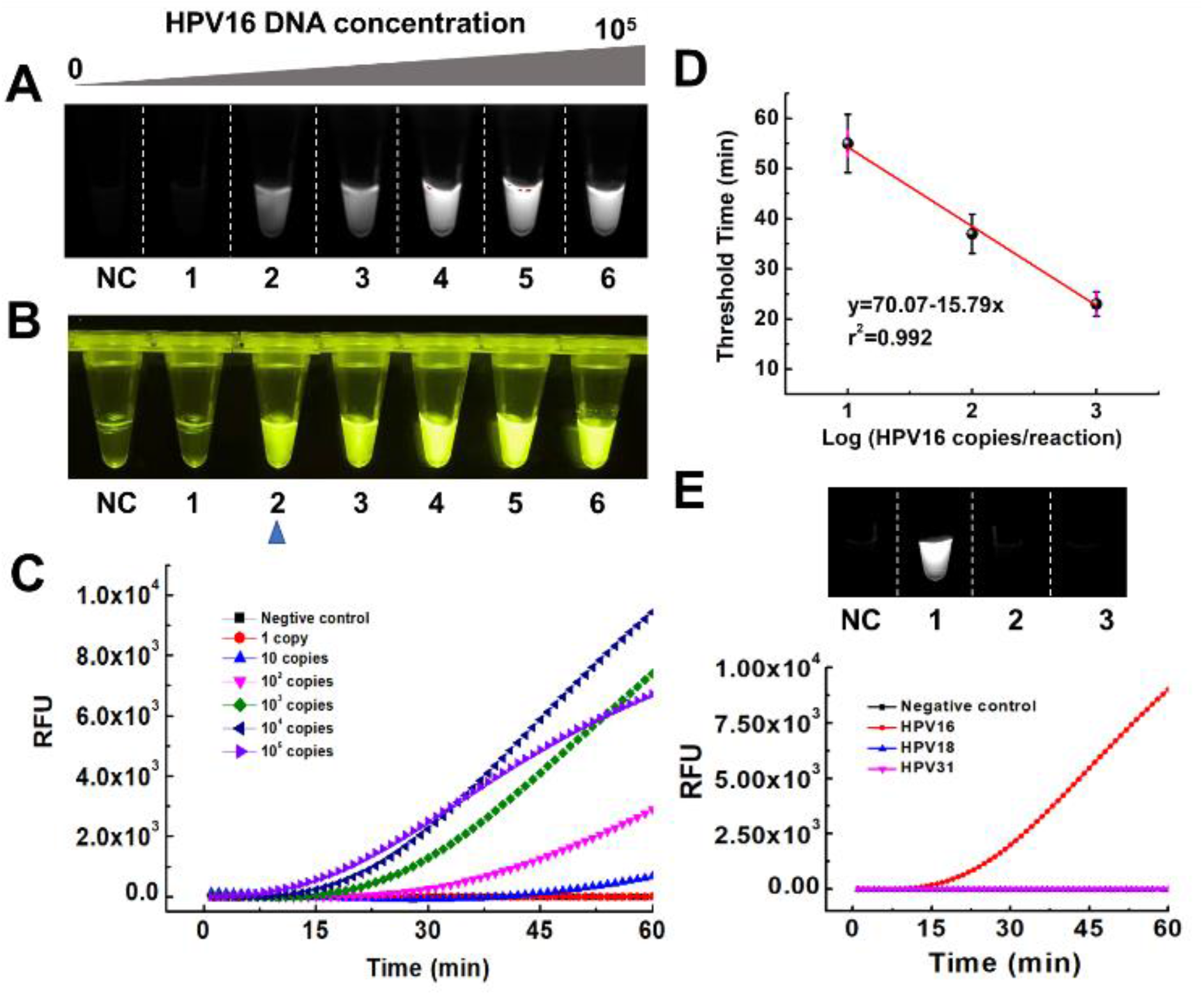
Quantitative detection of HPV DNA using one-pot RPA/CRISPR-Cas12a in the DAMR system. A-D. Ten-fold serial dilution of HPV 16 DNA (0, 1, 10, 10^2^, 10^3^, 10^4^, 10^5^ copies/reaction) was added into 20 μL RPA reaction bottom phase (with 10% sucrose) in the DAMR system and incubated at 37 °C for 1h. A. Endpoint fluorescent image taken by ChemiDocTM MP Imaging System. B. Endpoint fluorescent image taken by smartphone camera under blue light. C. Real-time fluorescent signal monitored by PCR machine. D. Linear relationship between the threshold time and HPV16 concentration (copies/reaction). (n=3) E. The selective detection of 10^4^ copies HPV 16 DNA over HPV18 and HPV31 using RPA/CRISPR-Cas12a detection in the DAMR system.

### Detection of high-risk genotype HPV16 and HPV18 in clinical samples

High-risk genotype HPV16 and HPV18 are responsible for 70% cervical cancer cases.^18, 19^ To evaluate the feasibility of our DAMR system for HPV clinical testing, DNA extracted from clinical human swab samples was first detected by the one-pot RPA/CRISPR-Cas12a in the DAMR system. As shown in Fig. 4A, HPV16 and HPV18 DNA were accurately identified by our DAMR within one hour and classified into four groups according to the threshold time difference, which was comparable with that of the conventional qPCR method.^20^ Next, we tested multiplexed detection ability of our DAMR system to simultaneously distinguish HPV 16 and 18, which is still a challenge in current CRISPR-based detection.^21^ We added both HPV16 and 18 primers in the RPA bottom phase and specific HPV16 or 18 crRNA in the top phase. As shown in Fig. 4B, HPV16/18 can be accurately identified and distinguished from each other. To further demonstrate the feasibility of our DAMR system for high-throughput multiplex detecion at point of care, we designed and fabricated a 3D-printed microfluidic device that can be inserted into the well of 96-well microplate (Fig.4C and S7). We used black polylactic acid (PLA) filament to fabricate 3D printed microfluidic devices because it showed low auto-fluorescence (Fig. S7A-D). As shown in Fig. S7E, three separated chambers of the 3D-printed devices did not influence each other, which should contribute to low diffusivity of the RPA bottom phase and the specific design of the insert 3D-printed device (Fig. 4C). As shown in Fig. 4D, both HPV16 and HPV18 primers were added into the RPA bottom phase and specific crRNA of HPV 16 or 18 were, respectively, added to the top phase in the respective chambers of the 3D-printed device. As shown in Fig. 4E, target HPV DNAs from clinical swab samples (Sample No.1, 11, 12 and 15) was successfully detected and identified in the 3D printed microfluidic devices after one-hour incubation at 37 °C. Therefore, all these results confirm that the DAMR system is able to achieve multiplexed detection of high-risk genotype HPV16 and HPV18 from clinical samples. Especially, by coupling with microfluidic technology, the DAMR system shows great potential for high-throughput CRISPR-based multiplex detection.

**Figure 4.**
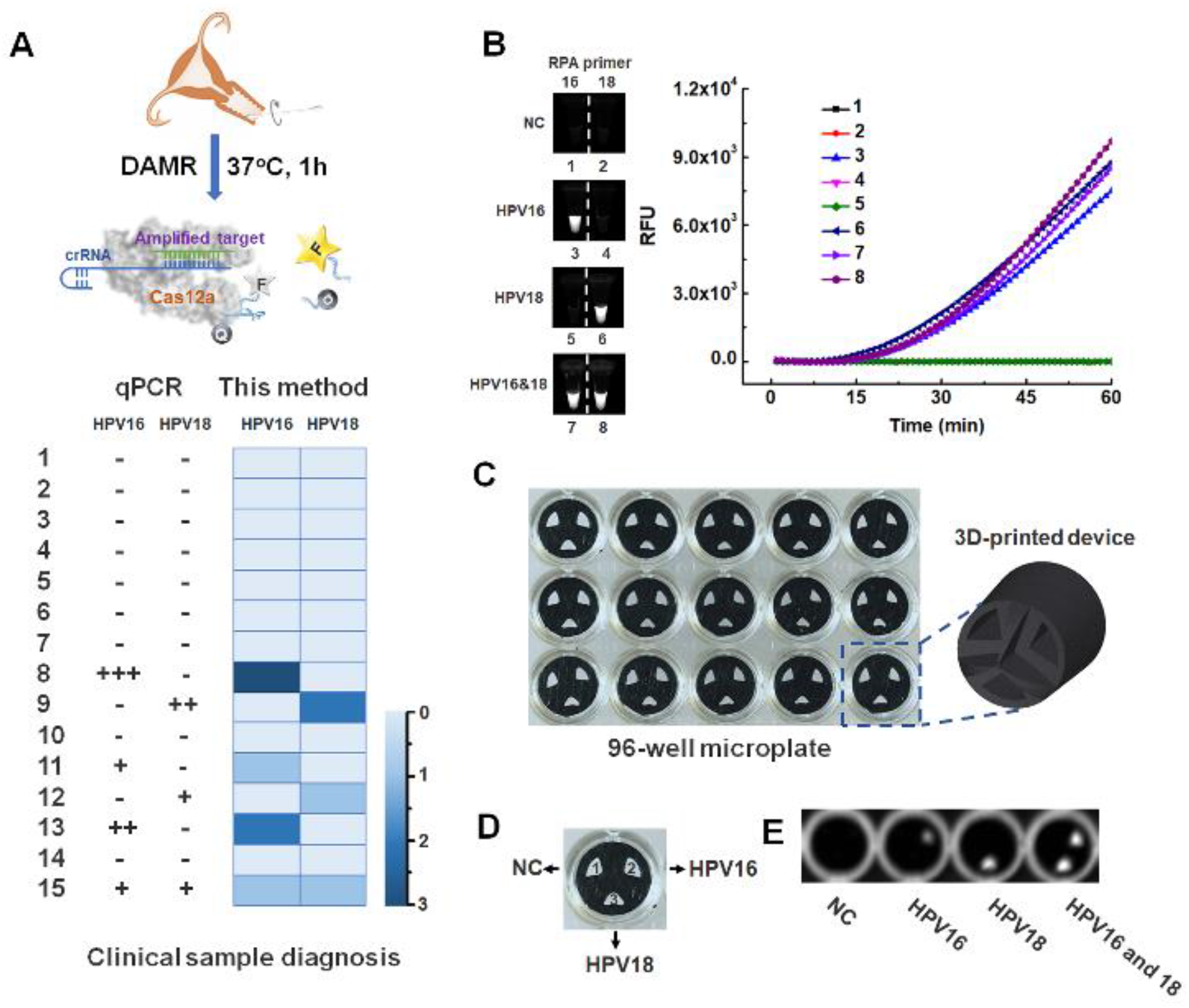
Multiple HPV DNA detection in clinical human swab sample. A. Top: Schematic outlining of the detection of HPV DNA from human swab samples by RPA/CRISPR-Cas12a assay in the DAMR system. Bottom: HPV 16/18 detection results of 15 clinical samples using qPCR (left) and RPA/CRISPR-Cas12a assay in the DAMR system (right), respectively. B. HPV16/18 detection in separate PCR tubes by RPA/CRISPR-Cas12a in the DAMR system. C. 3D-printed microfluidic device in 96-well microplate for multiplexed detection of HPV DNA in the DAMR system. D. Three chambers of 3D-printed device with or without specific crRNA (1: without crRNA, 2: with HPV16 crRNA and 3: with HPV18 crRNA). E. Endpoint fluorescent image of multiplex detection of HPV DNA with 3D printed device in the DAMR system.

### Detection of spiked HPV16 in untreated human plasma sample

Previous studies have shown that there are many inhibitors in human samples (e.g., plasma), such as immunoglobulin G, haemoglobin, lactoferrin and anticoagulant (e.g., EDTA), which can inhibit nucleic acid amplification detection (e.g., RPA, LAMP or PCR).^22–24^ It is necessary to extract and purify nucleic acids from human samples before enzymatic amplification, which is typically time consuming and tedious. To verify high tolerance of our DAMR system to the inhibitors, we directly detected HPV DNA spiked in untreated human plasma sample by using three-phase system. As shown in Fig. 5A, the three-phase system consists of: i) 10 μL plasma sample with 10^4^ copies of HPV16 DNA and 40% sucrose (bottom phase), ii) 20 μL RPA reaction solution with 10% sucrose (middle phase), and iii) 10 μL CRISPR-Cas12a solution (top phase). After one-hour incubation at 37 °C, significant fluorescent signals were detected in the three-phase DAMR system under an optimized sucrose concentration (Fig. 5B and C). In contrast, no fluorescent signal was observed in two-phase system in which the bottom phase and middle phase were mixed as one phase, which confirmed that the DAMR system minimized the effect of the inhibitors and was able to directly work raw human sample. To further evaluate the versatility of our DAMR system, we adapted it to detect HPV DNA in untreated plasma by the LAMP and PCR method, respectively (Fig. 5D). As shown in Fig. 5E and 5F, LAMP assay in our DAMR system worked well with raw plasma sample but qPCR method not, which may result from the rapid convection diffusion of the inhibitors in sample during PCR thermal cycling process. Therefore, the DAMR system provides a novel strategy to directly detect target DNA in clinical human samples without complicated sample pre-treatment.

**Figure 5.**
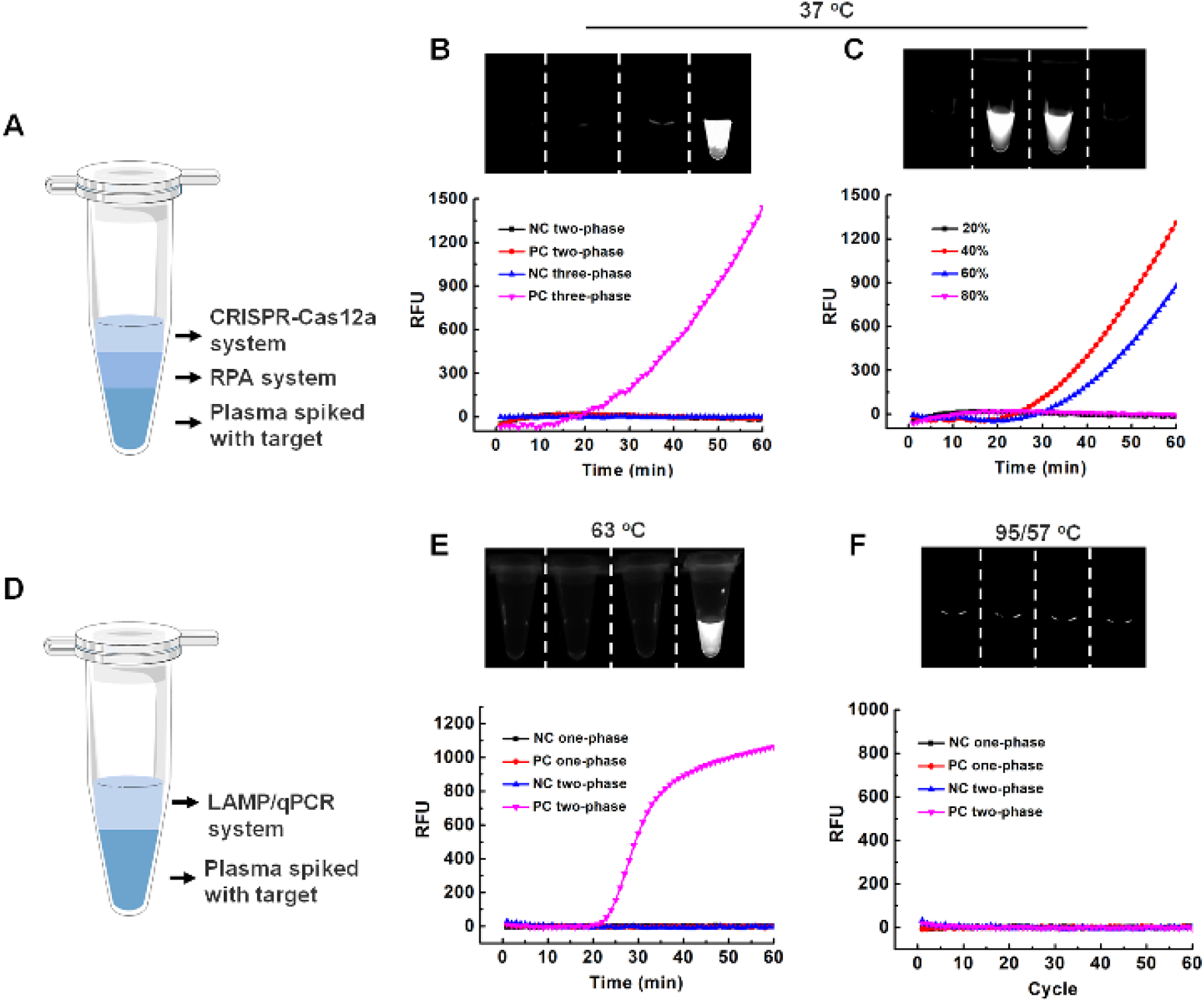
Direct DNA detection in untreated plasma sample in the DAMR system. A. Schematic illustration of three-phase system (plasma/RPA/CRISPR-Cas12a) for direct DNA detection using untreated plasma sample. B. Comparison of three-phase system and two-phase system for detection of HPV16 DNA in the untreated plasma. C. Optimization of sucrose concentration in the plasma bottom phase. D. Schematic illustration of two-phase system for DNA detection using untreated plasma samples by LAMP or qPCR method. E. Comparison of LAMP-based two-phase system and one-phase system for DNA detection in plasma samples. F. Comparison of PCR-based two-phase system and one-phase system for detection of DNA in plasma samples.

In summary, we developed a novel aqueous dynamic multiphase reaction system for simple, rapid, sensitive and quantitative detection of nucleic acid by using one-pot RPA/CRISPR-Cas12a assay. By combining with a 3D-printed microfluidic device, multiplex detection of HPV16 and 18 from clinical human swab samples was achieved using the DAMR system with excellent specificity and reliability. Additionally, we demonstrated that the our DAMR system has high tolerance to the inhibitors, enabling direct detection of nucleic acid in raw human samples (e.g., human plasma) at the point of care without need for complicated sample pre-treatment. We envision that such simple, robust, multiplexed, quantitative, CRISPR-based detection platform is compatible with various nucleic acid biomarkers and has a great potential for POC diagnosis and disease prevention.

## Methods

### Dynamic aqueous multiphase reaction (DAMR) system

The DAMR system was established using various concentrations of sucrose solution due to its good biocompatibility. Stock sucrose solutions at high concentrations were prepared in Milli-Q water. To investigate the dynamic diffusion of the DAMR system (Fig. S1A), we added equal volumes (15 μL) of sucrose solution (from 10% to 50%, w/w) and water into tubes and incubated at 37 °C for different times (0, 0.5 and 2h). In addition, sucrose solution from 10% to 30% were utilized to form multi-phase systems (Fig. S1B).

### One-pot RPA/CRISPR-Cas12a detection in the DAMR system

In the DAMR system, RPA-based target amplification and CRISPR-Cas12a based fluorescent detection was connected through dynamic diffusion. TwistAmp Basic reagent for the RPA reaction was purchased from TwistDx. High-density RPA reaction solution (bottom phase) was mixed by 0.48 μM forward and reverse primer, 14 mM magnesium acetate, targets and sucrose in 1× rehydration buffer. Low-density CRISPR-Cas12a detection solution (top phase) was mixed by 100 nM Cas12a, 50 nM ssDNA-FQ reporter, and 62.5 nM crRNA (Integrated DNA Technologies) in 1X cleavage buffer (100 mM KCl, 20 mM Tris-HCl (pH 7.8), 5 mM MgCl_2_, 50 μg mL^−1^ heparin, 1% (v/v) glycerol and 1 mM DTT). The DAMR systtem for one-pot RPA/CRISPR-Cas12a detection was incubated in PCR tubes at 37 °C and the real-time fluorescence signal was monitored by CFX96 Touch™ Real-Time PCR Detection System (Bio-Rad). After reaction, the endpoint fluorescent images were taken by ChemiDocTM MP Imaging System and the fluorescence intensity was analyzed by Image J. The bright-green images were taken directly by the smartphone camera under blue light.

### Optimization of RPA/CRISPR-Cas12a detection in the DAMR system

The dynamic diffusion between two phases is crucial to connect incompatible but correlative reactions, which depends on sucrose concentration and the volume ratio of RPA bottom phase and CRISPR-Cas12a top phase. To optimize the DAMR system, 20 μL RPA reaction solution with 10^4^ copies HPV16 DNA and different concentration of sucrose (0%, 5%, 10%, 20%, 30%, 40%, and 50%) was used as the bottom phase and 10 μL CRISPR-Cas12a detection solution was used as the top phase for one-pot RPA/ CRISPR-Cas12a detection at 37 °C. To optimize the volume ratio of RPA bottom phase (10% sucrose) and CRISPR-Cas12a top phase, different volume ratios (1: 1:5, 1:2, 1:1, 2:1, 5:1) was investigated under the same reaction condition.

### Sensitivity and specificity

To investigate the sensitivity of the DAMR system for one-pot RPA/CRISPR-Cas12a detection, ten-fold serial dilutions of HPV 16 or 18 DNA from 0 to 10^5^ copies were added into 20 μL RPA reaction phase with 10% sucrose with specific forward and reverse primers. HPV 16 or 18 specific crRNA was added in 10 μL CRISPR-Cas12a reaction top phase.^8^ The DAMR system was incubated in PCR tubes at 37 °C for 1h. To investigate the sensitivity of “two-step” method with seperate target amplification and detection steps, ten-fold serial dilutions of HPV 16 or 18 DNA from 0 to 10^5^ copies were firstly amplified in 20 μL RPA reaction at 37 °C for 15 min. Then 5 μL DNA amplicons from RPA reaction was added into 15 μL CRISPR-Cas12a reaction solution and incubaed at 37 °C for another 60 min. To investigate the specificity of DAMR system for one-pot RPA/CRISPR-Cas12a detection, 10^4^ copies HPV 16, 18 and 31 DNA was added into 20 μL RPA reaction bottom phase with 10% sucrose and HPV16 or 18 specific primers, respectively. HPV 16 or 18 crRNA was added into 10 μL CRISPR-Cas12a reaction solution as the top phase. The DAMR system was incubated at 37 °C in PCR tubes on CFX96 Touch™ Real-Time PCR Detection System for 1 hour.

### Multiplexed detection in PCR tubes and on 3D-printed microfluidic device in microplate

To investigated the multiplex detection ability of DARM system, 0.48 μM HPV 16 and HPV 18 RPA primers were mixed in 20 μL RPA reaction solution with 10% sucrose as the bottom phase. Specific HPV16 or HPV18 crRNA was added into 10 μL CRISPR-Cas12a reaction as the top phase. The DAMR system in PCR tubes was incubated at 37 °C for 1h to achieve multiplexed detection.

To realize high-throughput multiplexed detection, a microfluidic device with three respective chambers was fabricated by 3D-printing and was inserted into the wells of 96-well microplate. 0.48 μM HPV 16 and HPV 18 RPA primers were mixed into 70 μL RPA reaction as the bottom phase and added to the wells of the 96-well microplate. 3D-printed device was inserted into the wells and the respective chamber was added with 10 μL CRISPR-Cas12a reaction solution without crRNA, with HPV16 crRNA, and with HPV18 crRNA, respectively. After sealing with a Microseal^®^ B Adhesive sealer (Bio-Rad), the 96-well microplate was incubated at 37 °C for 1h and the end-point image was taken by ChemiDoc™ MP Imaging System (Bio-Rad).

### Human clinical swab sample collection and DNA purification

Clinical cervical swab samples were obtained from the Hospital of the University of Pennsylvania and approved by the ethics committee (IRB protocol #: 829760). To extract target nucleic acids from human clinical swab samples, 500 μL samples were centrifuged at 1000X g for 2 min to collect cell pellets. The cell pellets were suspended and washed by ddH_2_O for 3 times. Then cell pellets was resuspended and digested in 200 μL PBS solution with proteinase K. DNeasy Blood & Tissue Kit (Qiagen) was used for the extraction and purification of HPV DNA from the cell pellets. After purification, HPV DNA was directly used for detection or put in the freezer at −20 °C for strorage.

### Detection of HPV DNA in untreated human plasma sample

Human plasma samples were purchased from Innovative Research. In our three-phase system, 10^4^ copies of HPV16 DNA was added into 10 μL plasma sample with 40% sucrose as the bottom phase. 20 μL RPA reaction solution with 10% sucrose was used as the middle phase and 10 μL CRISPR-Cas12a reaction solution as the top phase. For comparison, we prepared two-phase system in which above-mentioned botton phase and middle phase were mixed togther as one phase. To optimize the sucrose concentration in the bottom plasma phase of our three-phase system, different concentration sucrose solutions (20%, 40%, 60% and 80%) were investigated.

In addition, target DNA in untreated human plasma samples were detected by LAMP and PCR by using our DAMR system. 10^4^ copies of HPV16 DNA was added into 10 μL plasma sample with 40% sucrose as the bottom phase and 20 μL LAMP or PCR solution was used as the top phase. For comparison, 10 μL plasma sample spiked with 10^4^ copies of HPV16 DNA was mixed to 20 μL LAMP or PCR solution as one phase. The LAMP reaction was incubated in PCR tubes at 63 °C for 1h.^25^ The PCR reaction was included at 95°C for 10 min, followed by a two-step cycle of 15 s at 95 °C and 1 min at 57 °C for 60 cycles.^20^

## Acknowledgements

The work was supported, in part, by R01EB023607, R01CA214072, and R21TW010625.

## References

1. Y. Du and S. Dong, Analytical chemistry, 2016, 89, 189–215.

2. Y. Li, S. Li, J. Wang and G. Liu, Trends in biotechnology, 2019.

3. R. K. Daher, G. Stewart, M. Boissinot and M. G. Bergeron, Clinical chemistry, 2016, 62, 947–958.

4. T. Notomi, H. Okayama, H. Masubuchi, T. Yonekawa, K. Watanabe, N. Amino and T. Hase, Nucleic acids research, 2000, 28, e63–e63.

5. R. Einspanier and A. Plath, in Molecular Biomethods Handbook, Springer, 1998, pp. 51–57.

6. H. Wang, M. La Russa and L. S. Qi, Annual review of biochemistry, 2016, 85, 227–264.

7. X. Wang, E. Xiong, T. Tian, M. Cheng, W. Lin, H. Wang, G. Zhang, J. Sun and X. Zhou, ACS nano, 2020.

8. J. S. Chen, E. Ma, L. B. Harrington, M. Da Costa, X. Tian, J. M. Palefsky and J. A. Doudna, Science, 2018, 360, 436–439.

9. J. S. Gootenberg, O. O. Abudayyeh, M. J. Kellner, J. Joung, J. J. Collins and F. Zhang, Science, 2018, 360, 439–444.

10. C. Myhrvold, C. A. Freije, J. S. Gootenberg, O. O. Abudayyeh, H. C. Metsky, A. F. Durbin, M. J. Kellner, A. L. Tan, L. M. Paul and L. A. Parham, Science, 2018, 360, 444–448.

11. J. S. Gootenberg, O. O. Abudayyeh, J. W. Lee, P. Essletzbichler, A. J. Dy, J. Joung, V. Verdine, N. Donghia, N. M. Daringer and C. A. Freije, Science, 2017, 356, 438–442.

12. S.-Y. Li, Q.-X. Cheng, J.-M. Wang, X.-Y. Li, Z.-L. Zhang, S. Gao, R.-B. Cao, G.-P. Zhao and J. Wang, Cell discovery, 2018, 4, 20.

13. B. Wang, R. Wang, D. Wang, J. Wu, J. Li, J. Wang, H. Liu and Y. Wang, Analytical chemistry, 2019, 91, 12156–12161.

14. M. K. Brakke, Journal of the American Chemical Society, 1951, 73, 1847–1848.

15. V. Molinero, T. Çaǧın and W. A. Goddard Iii, Chemical physics letters, 2003, 377, 469–474.

16. J. Hindmarsh, A. Russell and X. D. Chen, Journal of crystal growth, 2005, 285, 236–248.

17. H. Sales-Ortells, G. Agostini and G. Medema, Environmental science & technology, 2015, 49, 6943–6952.

18. M. Schiffman, P. E. Castle, J. Jeronimo, A. C. Rodriguez and S. Wacholder, The Lancet, 2007, 370, 890–907.

19. G. S. Ogilvie, D. van Niekerk, M. Krajden, L. W. Smith, D. Cook, L. Gondara, K. Ceballos, D. Quinlan, M. Lee and R. E. Martin, JAMA, 2018, 320, 43–52.

20. M. Moberg, I. Gustavsson and U. Gyllensten, Journal of clinical microbiology, 2003, 41, 3221–3228.

21. Y. Li, L. Liu and G. Liu, Trends in biotechnology, 2019, 37, 792–795.

22. W. A. Al-Soud and P. Rådström, Journal of clinical microbiology, 2001, 39, 485–493.

23. C. Schrader, A. Schielke, L. Ellerbroek and R. Johne, Journal of applied microbiology, 2012, 113, 1014–1026.

24. H. Kaneko, T. Kawana, E. Fukushima and T. Suzutani, Journal of biochemical and biophysical methods, 2007, 70, 499–501.

25. K. Yin, V. Pandian, K. Kadimisetty, C. Ruiz, K. Cooper, J. You and C. Liu, Theranostics, 2019, 9, 2637.

