## Supporting information for "Dynamic aqueous multiphase reaction system for simple, sensitive and quantitative one-pot CRISPR-Cas12a based molecular diagnosis"

Dr. Changchun Liu

Department of Biomedical Engineering
University of Connecticut Health Center
263 Farmington Avenue, Farmington, CT 06030, USA

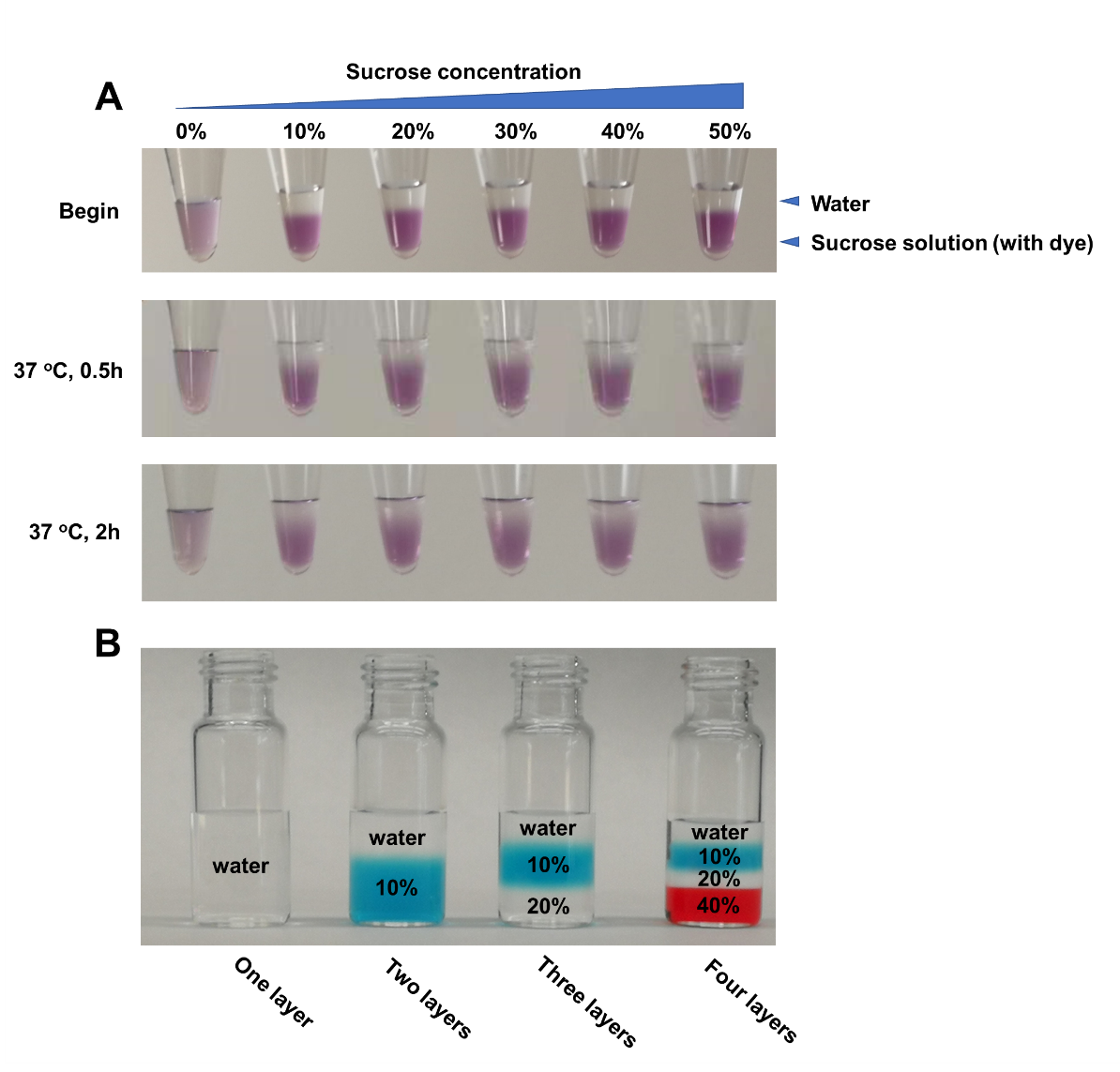

**Fig. S1 | Dynamic aqueous multiphase reaction (DAMR) system.** A. Photo of the DAMR system with various sucrose concentration from 10% to 50% (w/w) at different incubation times (0, 0.5 and 2h). B. Photo of the DAMR system with various phases by adding sucrose with different concentrations (from 10% to 40%, w/w). Dye was added to indicate phase boundaries.

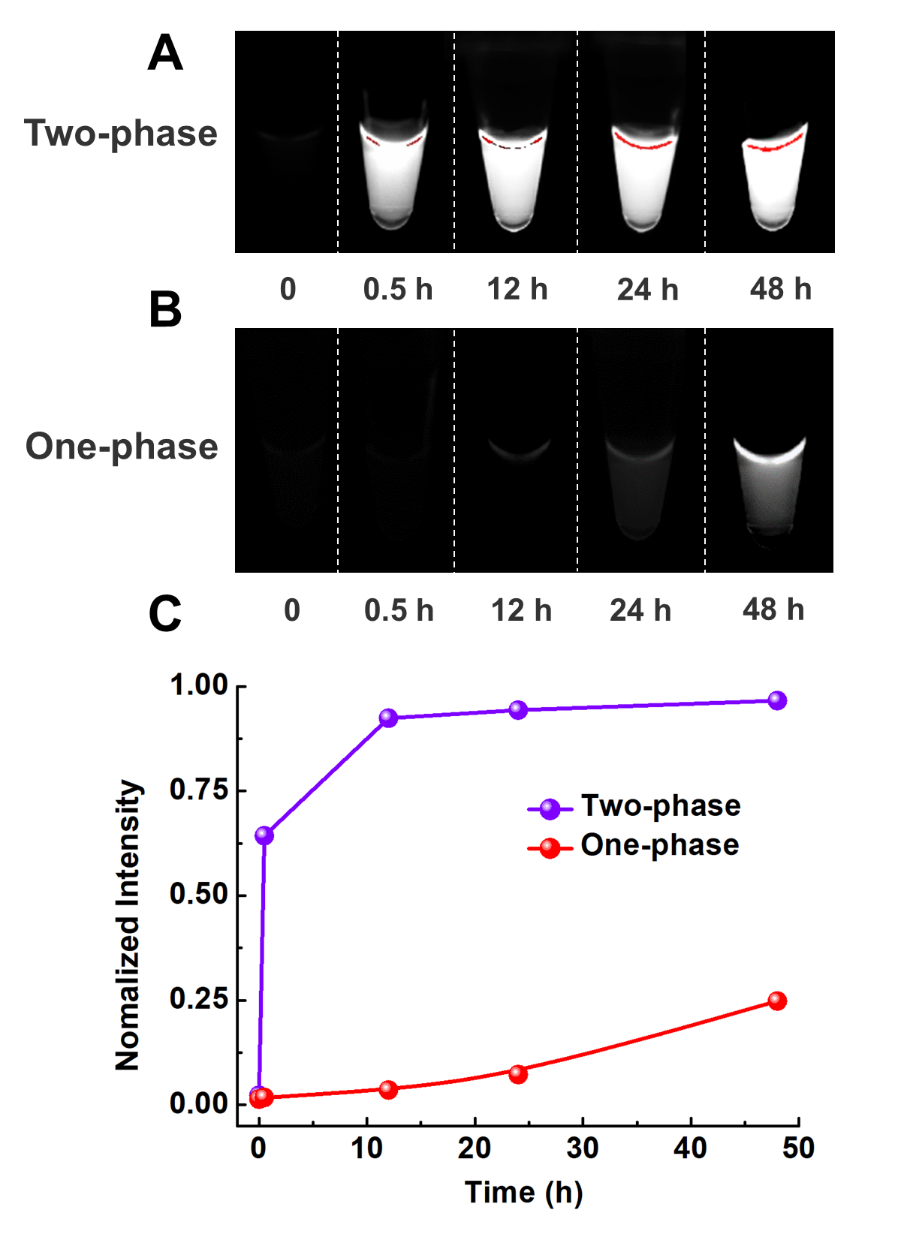

**Fig. S2 | Comparison of one-pot RPA/CRISPR-Cas12a detection in the DAMR system and one-phase system.** A. Fluorescent images of one-pot RPA/CRISPR-Cas12a reaction in the DAMR system at different incubation times. B. Fluorescent images of one-pot RPA/CRISPR-Cas12a reaction in the one-phase system at different incubation times. C. Fluorescent monitoring of one-pot RPA/CRISPR-Cas12a reaction at different incubation times in the DAMR and one-phase system, respectively.

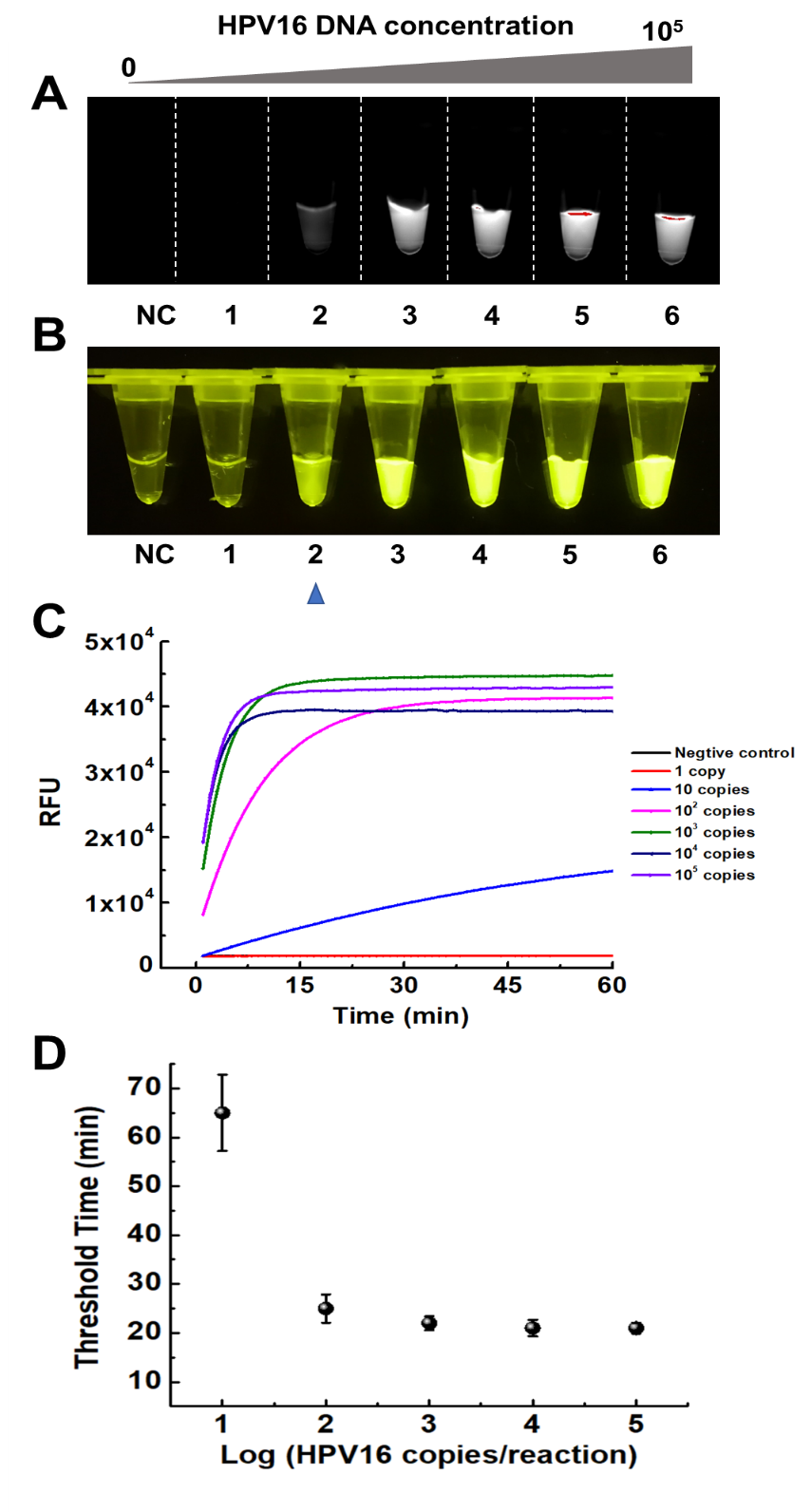

**Fig. S3 | HPV 16 DNA detection using "two-step" method.** Ten-fold serial dilution of HPV 16 DNA (0, 1, 10, 10^2^, 10^3^, 10^4^, 10^5^ copies/reaction) was first added into 20 μL RPA reaction and incubated at 37 ^o^C for 15 min. Then, 5 μL RPA reaction solution was added into 15 μL CRISPR-Cas12a detection solution and incubated at 37 ^o^C for another 60 min. A. Endpoint fluorescent image taken by ChemiDocTM MP Imaging System. B. Endpoint fluorescent image taken by smartphone camera under blue light. C. Real-time fluorescence curves were monitored by real-time PCR machine. D. Threshold time (min) of the detection of different concentration HPV16. (n=3).

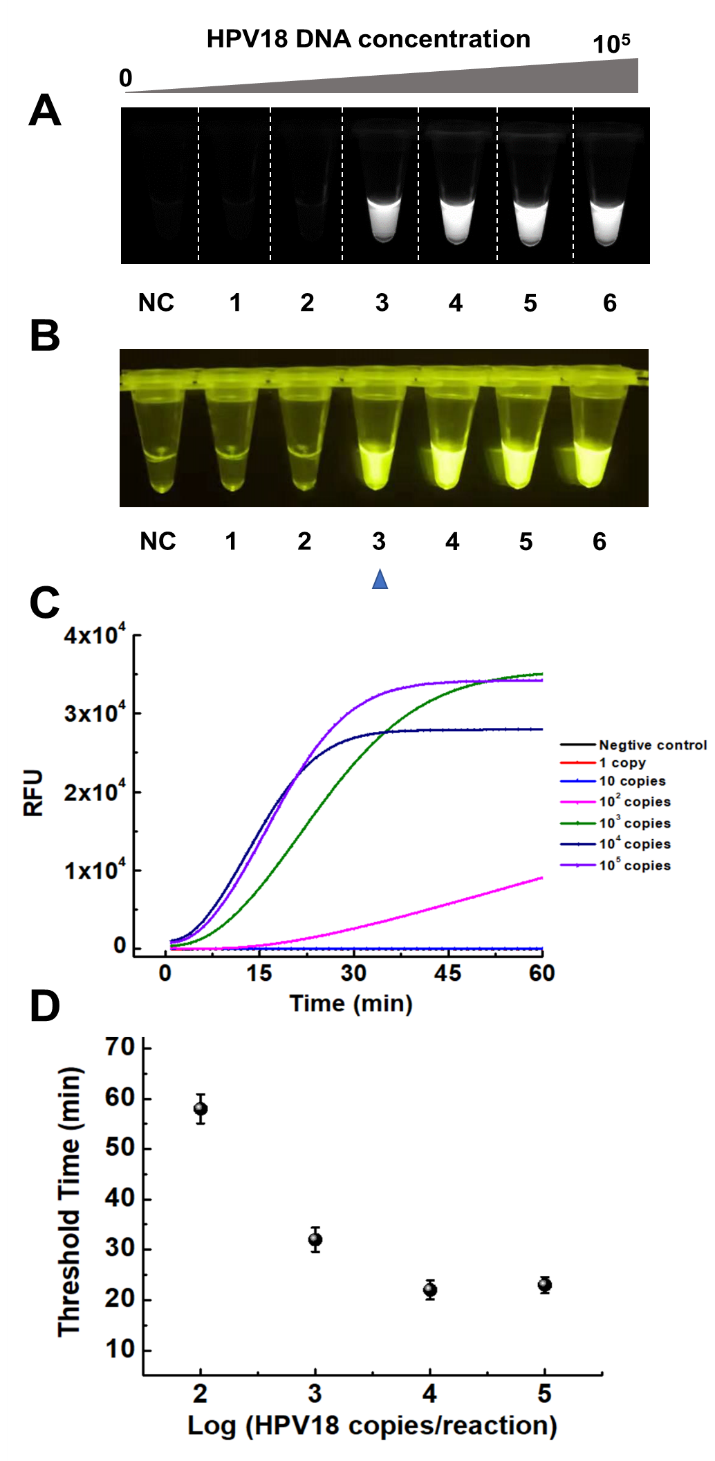

**Fig. S4 | HPV 18 DNA detection** **using "two-step" method.** Ten-fold serial dilution of HPV 18 DNA (0, 1, 10, 10^2^, 10^3^, 10^4^, 10^5^ copies/reaction) was first added into 20 μL RPA reaction and incubated at 37 ^o^C for 15 min. Then, 5 μL RPA reaction solution was added into 15 μL CRISPR-Cas12a detection solution and incubated at 37 ^o^C for another 60 min. A. Fluorescent image taken by ChemiDocTM MP Imaging System after 60-min incubation. B. Fluorescent image taken by smartphone camera in blue light transilluminator after 60-min incubation. C. Real-time fluorescence curves were recorded by real-time PCR machine. D. Threshold time (min) of the detection of different concentration HPV18. (n=3).

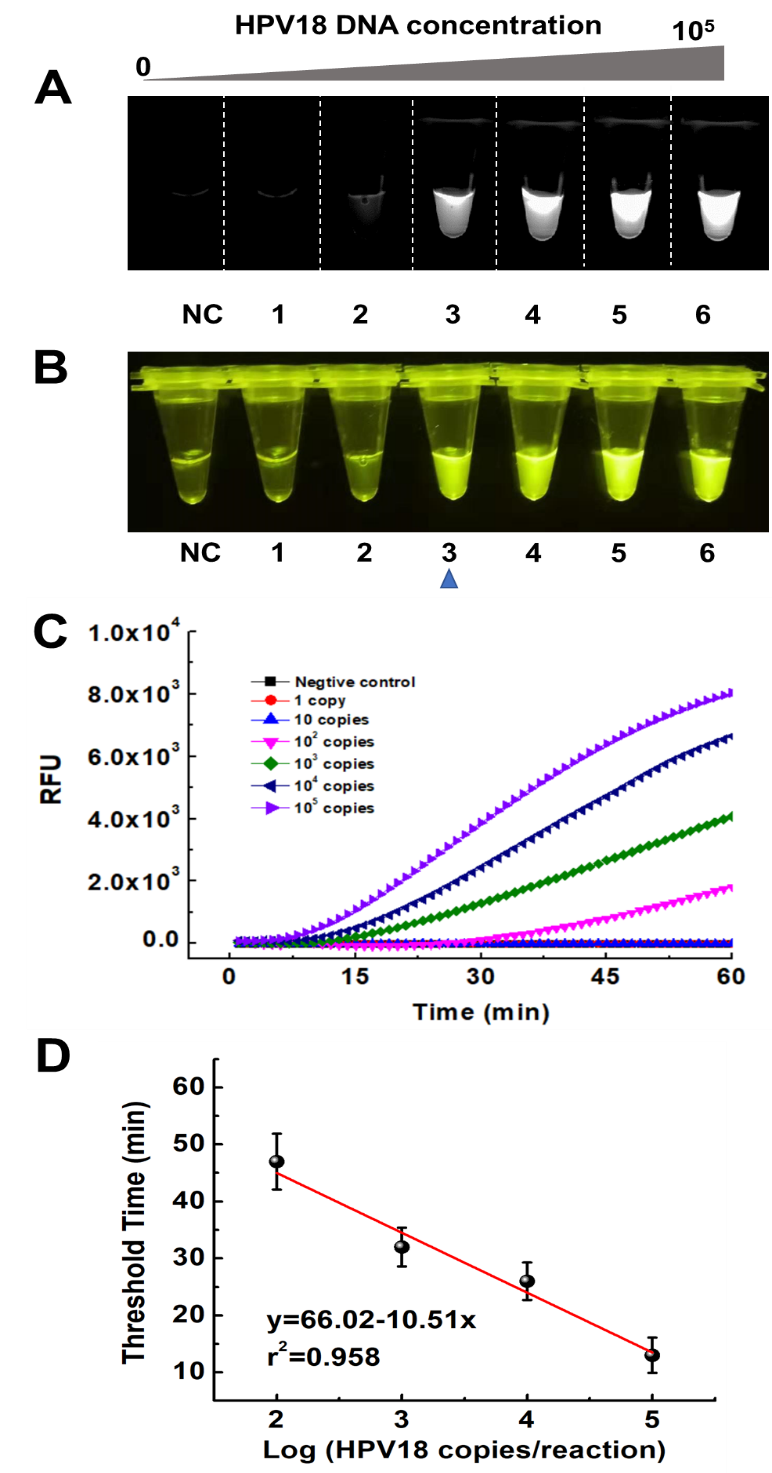

**Fig. S5 | HPV 18 DNA detection by one-pot RPA/CRISPR-Cas12a in the DAMR system.** Ten-fold serial dilution of HPV 18 DNA (0, 1, 10, 10^2^, 10^3^, 10^4^, 10^5^ copies/reaction) was added into 20 μL RPA reaction bottom phase (with 10% sucrose) in the DAMR system and incubated at 37 ^o^C for 1h. A. Endpoint fluorescent image taken by ChemiDocTM MP Imaging System. B. Endpoint fluorescent image taken by smartphone camera under blue light. C. Real-time fluorescent signal collected by PCR machine. D. Linear relationship between the threshold time and HPV18 concentration (copies/reaction). (n=3)

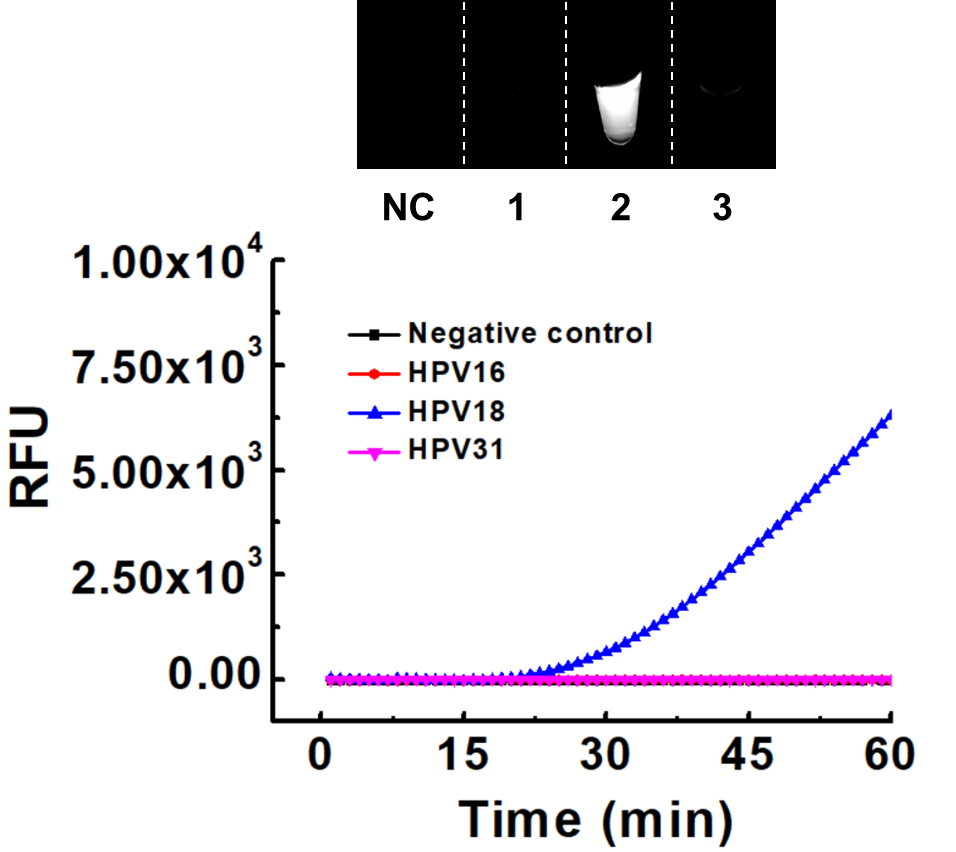

**Fig. S6 | Selective detection of HPV18 DNA by one-pot RPA/CRISPR-Cas12a in the DAMR system.** 10^4^ copies HPV 16, 18 and 31 DNA was added into 20 μL RPA reaction bottom phase (10% sucrose and HPV18 RPA primers) in the DAMR system and incubated at 37 ^o^C for 1h, respectively. Top: Endpoint fluorescent image taken by ChemiDocTM MP Imaging System. Bottom: real-time fluorescent signal curves were monitored by real-time PCR machine.

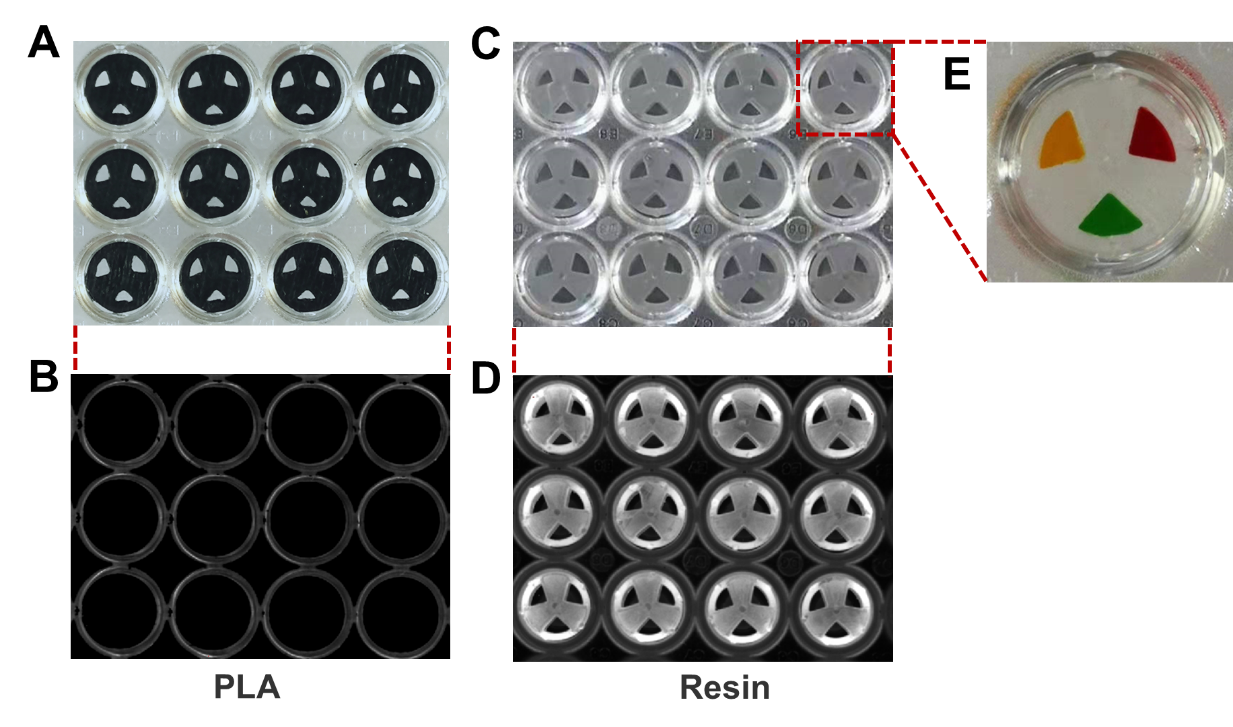

**Fig. S7 | 3D-printed microfluidic devices in 96-well microplate.** A-B. 3D-printed microfludic device printed by PLA material (A: optical image under bright view, B: fluorescent image taken by ChemiDocTM MP Imaging System). C-D. 3D-printed microfluidic device printed by clear resin material (C: optical image under bright view, D: fluorescent image taken by ChemiDocTM MP Imaging System). E. Photo of the 3D-printed microfludic device containing different dyes in three independent chambers after one-hour incubation at 37 ^o^C.

### Reference
